# Investigation of putative regulatory acetylation sites in Fas2p of *Saccharomyces cerevisiae*

**DOI:** 10.1101/430918

**Authors:** Alexandra Bergman, Leonie Wenning, Verena Siewers, Jens Nielsen

**Affiliations:** Department of Biology and Biological Engineering, Systems and Synthetic Biology, Chalmers University of Technology, Kemivägen 10, SE-412 96 Göteborg, Sweden; Novo Nordisk Foundation Center for Biosustainability, Chalmers University of Technology, Kemivägen 10, SE-412 96 Göteborg, Sweden; Novo Nordisk Foundation Center for Biosustainability, Technical University of Denmark, Kemitorvet, DK-2800 Kgs. Lyngby, Denmark

**Keywords:** *Saccharomyces cerevisiae*, fatty acid synthase, regulation, protein acetylation, residue replacement system

## Abstract

Yeast metabolism is highly regulated, in part via coordinated reprogramming of metabolism on a transcriptional level, for example in response to environmental changes. Furthermore, regulation occurs on the protein level via posttranslational modifications directly affecting enzymatic activity – a mode of regulation that has the benefit of being very fast in response to environmental changes. One group of posttranslational modification that has been suggested to have a high impact on regulation of metabolism are acetylations. Around 4000 distinct protein acetylation sites have been found in *Saccharomyces cerevisiae*, many of which are located in central metabolic enzymes. However, reports on the verification of regulatory roles of specific acetylation sites on these metabolic enzymes have yet to emerge. This study investigates putative regulatory acetylation sites on Fas2p, which in concert with Fas1p is responsible for cytosolic fatty acid (FA) biosynthesis in *S. cerevisiae*. Fas2p stands out as one of the most highly acetylated proteins in yeast and is located at a branchpoint of acetyl-CoA metabolism. The amino acids (AAs) glutamine (Q) and arginine (R) were introduced to mimic a constitutively acetylated or non-acetylatable state at three separate lysine sites (K) (K83, K173 and K1551) confirmed to be acetylated in two independent studies, either separately or simultaneously. The results suggest that the residue replacement system in the specific case interferes with the enzymatic function of the fatty acid synthase (FAS), as QQQ and RRR triple mutants both reduce the amount of secreted free fatty acids (FFAs) in a *faa1*Δ *faa4*Δ yeast deletion mutant. The K173Q substitution significantly decreased C16 FA species at the expense of C18 FAs, while no such change could be observed for the corresponding K173R modification.

## Introduction

Acetyl-coenzyme A (CoA) is a central metabolite in yeast, which results from the degradation of glucose via glycolysis in the cytosol, but also plays an important role in other catabolic and anabolic pathways in different organelles, including mitochondria, the nucleus and peroxisomes.

Cytosolic acetyl-CoA is formed in three steps from pyruvate, a product of glycolysis. The first step is the formation of acetaldehyde by deacetylation of pyruvate, catalyzed by a pyruvate decarboxylase (PDC). In yeast, three different PDCs (Pdc1p, Pdc5p and Pdc6p) exist, with Pdc1p being responsible for the majority of PDC activity in glucose-rich medium (Pronk et al., 1996). In general, high glucose concentrations promote the conversion of acetaldehyde towards ethanol by an alcohol dehydrogenase (Adh1p, Adh3p, Adh4p and Adh5p) and only a small fraction is converted to acetate by an aldehyde dehydrogenase (Ald2p, Ald3p, Ald6p) (Gombert et al., 2001; Heyland et al., 2009). The majority of acetate in the cytosol is produced by Ald6p in glucose rich medium (Dickinson, 1996; Meaden et al., 1997). Finally, acetyl-CoA is formed by the action of acetyl-CoA synthetase (ACS) under consumption of ATP. Yeast possesses two ACSs enzymes (Acs1p and Acs2p), with *ACS1* being repressed in glucose containing medium and Acs2p therefore being essential for acetyl-CoA formation under those conditions (van den Berg et al., 1996; Kratzer and Schüller, 1995; Starai and Escalante-Semerena, 2004; Takahashi et al., 2006). Acs1p was found to be localized in the cytosol, the nucleus and peroxisomes, whereas Acs2p was detected in the cytosol, the nucleus and perhaps the endoplasmic reticulum (Chen et al., 2012; Huh et al., 2003; Kals et al., 2005; Takahashi et al., 2006).

Since glucose represses the tricarboxylic acid (TCA) cycle and respiration, only a small amount of pyruvate is transported from the cytosol into mitochondria under high glucose conditions (Gombert et al., 2001; Heyland et al., 2009; Weinert et al., 2014). Mitochondrial acetyl-CoA is the product of the oxidative decarboxylation of pyruvate by the pyruvate dehydrogenase (PDH) complex in the mitochondrial matrix, whereas peroxisomal acetyl-CoA is the product of FA degradation via β-oxidation but is also synthesized from acetate. Studies with a heterologous *Salmonella enterica* ACS in a yeast strain deficient in Acs2p indicate that acetyl-CoA in the cytosol and the nucleus form a single nucleocytosolic acetyl-CoA pool and that acetyl-CoA can move freely through the nuclear pores, but that mitochondrial acetyl-CoA is biochemically separated (Takahashi et al., 2006). Because of that, different transport mechanisms exist for acetyl-CoA. One of these mechanisms is based on the free movement of acetate between different compartments in the cell. In mitochondria, the hydrolysis of acetyl-CoA to acetate and CoA is catalyzed by the glucose-repressed acetyl-CoA hydrolase Ach1p (Buu et al., 2003; Lee et al., 1989; Lee et al., 1990). Two other reactions catalyzed by Ach1p include the CoA transfer from succinyl-CoA to acetate to form acetyl-CoA and succinate (Fleck and Brock, 2009) and the potential transfer of CoA from acetyl-CoA to succinate to form acetate and succinyl-CoA under glucose derepressed conditions in a PDC negative *S. cerevisiae* strain also containing a deletion in *MTH1* (Chen et al., 2015). A second option for acetyl-CoA transfer is via acetyl-carnitine which can be formed in the cytosol, the mitochondria and the peroxisomes from acetyl-CoA and carnitine, catalyzed by one of three carnitine acetyl-CoA transferases in *S. cerevisiae* (Elgersma et al., 1995; Schmalix and Bandlow, 1993; Swiegers et al., 2001). This way of transport is only active when carnitine is externally supplied though (van Roermund et al., 1999). The third option for acetyl-CoA transport is via the cytosolic/peroxisomal glyoxylate cycle. This pathway is essential for growth on C_2_ compounds (like acetate and ethanol), but also for degradation of FAs via β-oxidation, and produces succinate which can enter the TCA cycle (van Roermund et al., 1995).

As a central metabolite, acetyl-CoA is an important building block in many anabolic reactions, but it also plays a role in the posttranslational modification of proteins by serving as a substrate for lysine acetyltransferases (KATs), which catalyze the transfer of acetyl groups to the epsilon-amino groups of lysines (Lys, K). In this way, fluctuating concentrations of acetyl-CoA, which are connected to the metabolic condition of the cell, are translated into dynamic protein acetylations that regulate a range of cellular functions. Enzymes that remove acetylations from proteins are called lysine deacetylases (KDACs). So far, seven KATs and nine KDACs, respectively, have been identified in yeast. KATs are classified based on their cellular localization, with being either of the nuclear type (A-type) or the cytoplasmic type (B-type), whereas KDACs are phylogenetically divided into class I, class II and class III enzymes (also called sirtuins) (Galdieri et al., 2014). The KDACs of class I and II have similar catalytic domains, while sirtuins (class III) are NAD^+^ dependent (Blander and Guarente, 2004; Ekwall, 2005).

The acetylation of proteins is a widespread phenomenon in pro- and eukaryotic species and has been studied in bacteria, like *Escherichia coli*, but also in complex organisms like *Mus musculus* and *Homo sapiens* (Choudhary et al., 2014). In a *S. cerevisiae* strain carrying a deletion in the KDAC Rpd3p (*rpd3*Δ), over 4000 acetylation sites have been detected via high resolution mass spectrometry in batch grown cells harvested in the exponential phase (OD_600_ ~ 0.5) (Henriksen et al., 2012).

The degree of acetylation in the different cellular compartments of yeast is regulated by the level of acetyl-CoA, which is highest during exponential growth on glucose and lower during the diauxic shift, the ethanol phase and the following stationary phase. Current methods for acetyl-CoA measurements cannot distinguish between nucleocytosolic and mitochondrial acetyl-CoA. Therefore, the total cellular level of acetyl-CoA is usually used to assess nucleocytosolic levels (Cai et al., 2011; Ramaswamy et al., 2003; Sandmeier et al., 2002; Seker et al., 2005). A deletion of *PDA1*, encoding the E1 alpha subunit of the PDH complex, leads to a 33% decrease in cellular acetyl-CoA levels, which leads to the assumption that the highest concentration of acetyl-CoA can be found in the mitochondria. Since mitochondria only occupy 1–2% of the cellular volume in early log-phase under glucose excess, the concentration of acetyl-CoA in the mitochondria is about 20–30 fold higher than in the nucleocytosol (Cai et al., 2011; Uchida et al., 2011; Wagner and Payne, 2013; Weinert et al., 2014). This ratio might change when cells go through the diauxic shift into stationary phase. Moreover, so far it is not known if the levels of acetyl-CoA in the nucleocytosol and the mitochondria influence and control each other (Galdieri et al., 2014). Besides being catalyzed by KATs, acetylation of proteins can also occur spontaneously, especially in the mitochondria where the concentration of acetyl-CoA as well as the pH are higher than in the cytosol or the nucleus (Wagner and Payne, 2013; Weinert et al., 2014).

The nucleocytosolic acetyl-CoA pool is also influenced by *de novo* fatty acid (FA) biosynthesis in the cytosol which uses acetyl-CoA as a building block. The first rate limiting step in *de novo* FA biosynthesis is the formation of malonyl-CoA from acetyl-CoA catalyzed by an acetyl-CoA carboxylase (Acc1p) (Henry et al., 2012; Tehlivets et al., 2007). When the expression of *ACC1* is reduced, global histone acetylation is increased by an elevated level of acetyl-CoA in the nucleocytosol, which in turn leads to an altered transcriptional regulation (Galdieri and Vancura, 2012; Zhang et al., 2013).

Another factor regulating nucleocytosolic acetyl-CoA concentration is the Snf complex which is the budding yeast ortholog of mammalian AMP activated protein kinase (AMPK) (Carling et al., 1994; Woods et al., 1994; Zhang et al., 2013). The SNF1/AMPK pathway is highly conserved in eukaryotes and plays a role in energy sensing of cells as well as in regulating metabolism (Hardie, 2007; Hedbacker and Carlson, 2008). Snf1 plays a critical role in the transition from calorie rich to calorie restricted condition in aerobic environment and the reprogramming of metabolism from fermentation to respiration by controlling metabolic pathways involved in ATP production and consumption (Zhang et al., 2011). The SNF1 complex is composed of the catalytic α-subunit Snf1p, one of three different regulatory β- subunits (Siplp, Sip2p or Gal83p) and the stimulatory γ-subunit Snf4p (Jiang and Carlson, 1997). A protein phosphorylated, and thereby inactivated, by Snf1p is Acc1p (Witters and Watts, 1990; Woods et al., 1994). Therefore, the inactivation of Snf1p results in increased Acc1p activity and an increased conversion of acetyl-CoA to malonyl-CoA which reduces the pool of acetyl-CoA in the nucleocytosol and thereby leads to a decreased global histone acetylation. Moreover, *snf1*Δ mutants show a reduced fitness and reduced stress resistance (Zhang et al., 2013). One of the regulatory β-subunits of the SNF1 complex (Sip2p) is subject to acetylation (Lin et al., 2009). The level of acetylation is dependent on the acetyl-CoA concentration in the cell and is decreased in *snf1*Δ mutants (Galdieri and Vancura, 2012; Zhang et al., 2013). The acetylation of Sip2p leads to an increased interaction with Snf1p and thereby inhibition of Snf1p, which in turn leads to increased activity of Acc1p by decreased phosphorylation. On the other hand, decreased acetyl-CoA levels lead to hypoacetylation of Sip2p, and thereby increased activity of Snf1p, which in turn leads to increased phosphorylation and inhibition of Acc1p as well as decreased conversion of acetyl-CoA to malonyl-CoA (Lu et al., 2011). Acc1p is not only subject to posttranslational modifications, but is also regulated on the transcriptional level, with high nucleocytosolic concentrations of acetyl-CoA leading to increased histone acetylation at the promoter region of *ACC1*, and thereby to increased *ACC1* expression levels, which in turn translates to an increased conversion of acetyl-CoA to malonyl-CoA (Galdieri et al., 2014).

Other factors that probably influence the acetyl-CoA concentration are the level of ATP, since ACS enzymes are dependent on ATP, as well as the level of NAD^+^ or the ratio of NAD^+^/NADH, since the deacetylases of the sirtuin type are dependent on this cofactor (Guarente, 2011; Starai et al., 2004).

The most studied proteins in terms of protein acetylation are histones, which belong to the DNA-packaging proteins. Four different histones (H2A, H2B, H3 and H4) form an octamer, and together with 147 bp of DNA a nucleosome, which are the building blocks of chromatin. The acetylation of histones, which occurs at all four core histones at their amino terminal tails, enables the control of aging, cell cycle progression, DNA repair, replication and transcription. In general, increased acetylation of histones located at promoter regions results in increased transcription (Galdieri et al., 2014). One of the KATs, catalyzing various acetylations of histone H3 in yeast, is Gcn5p. (Burgess et al., 2010; Govind et al., 2007). It is a homolog of the human GCN5, is involved in regulation of gluconeogenesis and interacts with the deacetylase Sir2p (Dominy et al., 2012). Sir2p is a KDAC of the sirtuin type, belongs to the SIR complex, and is responsible for the deacetylation of histone H4 at K16 in yeast and thereby for the assembly of heterochromatin. Since Sir2p deacetylation activity is dependent on NAD+, it functions as a redox sensor. Moreover, it activates respiration and directs the flux towards the TCA cycle by an so far not known mechanism (Guarente, 2011; Lin et al., 2000; Lin et al., 2002). Sir2p is the human homolog of SIRT1 and has been studied extensively for its role in aging, genome stability and protein quality control (Lin et al., 2002; Liu et al., 2010).

Besides histones, also several other proteins involved in cell cycle progression, cytokinesis, metabolism, RNA processing, stress response and transcription have been identified as targets for acetylation in yeast (Duffy et al., 2012; Lin et al., 2009; Weinert et al., 2014) as well as other organisms including bacteria (Starai et al., 2002; Starai et al., 2003; Starai et al., 2004) and mammals (Hallows et al., 2006; Lin et al., 2016; Schwer et al., 2006; Wang et al., 2014). In yeast, a large proportion of metabolic enzymes involved in glycolysis, gluconeogenesis and amino acid (AA) metabolism were found to be acetylated (Henriksen et al., 2012). Many of these enzymes are highly conserved and have been reported to be acetylated in other organisms, suggestive of that regulation by acetylation has a conserved role in cellular metabolism. One example for an acetylated metabolic protein in yeast is Pck1p, encoding phosphoenolpyruvate carboxykinase, which catalyzes the conversion of oxaloacetate to phosphoenolpyruvate in gluconeogenesis. In this case, acetylation of K514 is crucial for enzymatic activity and for the ability of yeast cells to grow on non-fermentable carbon sources (Lin et al., 2009). Another prominent example is the regulation of bacterial and mammalian ACSs by acetylation. The acetylation of a conserved lysine residue in *Salmonella enterica* ACS leads to an almost complete inactivation of the enzyme, whereas deacetylation can reactivate the enzyme. Direct evidence for the role of acetylation on the activity of *S. cerevisiae* ACS is lacking so far (Hallows et al., 2006; Schwer et al., 2006; Starai et al., 2002; Starai et al., 2003; Starai et al., 2004). In case of the human glucose-6-phosphate dehydrogenase (G6PD), which is a key enzyme in the pentose phosphate pathway (PPP) that plays an essential role in the oxidative stress response by producing NADPH, the acetylation of K403 leads to inability of the enzyme to form active dimers and therefore to a complete loss of enzyme activity (Wang et al., 2014). The acetylation of human fatty acid synthase (FAS) destabilizes the enzyme by promoting its degradation via the ubiquitin-proteasome pathway which leads to a decreased *de novo* lipogenesis as well as tumor cell growth. This makes acetylation of human FAS a potential anticancer drug target (Lin et al., 2016).

Acetylation sites have also been detected in the FAS of *S. cerevisiae* in two independent studies (Henriksen et al., 2012; Kumar et al., unpublished). The FAS of *S. cerevisiae* consists of the β-subunit Fas1p and the α-subunit Fas2p which together form a hexameric α6β6 complex. This enzyme complex is responsible for the synthesis of FAs in the cytosol of *S. cerevisiae* based on the precursor molecules acetyl-CoA and malonyl-CoA (Tehlivets et al., 2007). Fas1p and Fas2p were found to have the highest number of detected unique acetylation sites in *S. cerevisiae* (29 and 50 sites, respectively). When normalized to protein length, this represents an acetylation frequency of 1.41 and 2.65 detected acetylations/100 AAs in Fas1p and Fas2p, respectively. Therefore, Fas1p and Fas2p likely belong to the top 20% and top 10% of yeast proteins with the highest frequency of acetylation, respectively (Henriksen et al., 2012). Three of these lysine acetylation sites in Fas2p (K83, K173 and K1551) have been confirmed in a study by Kumar et al. in which six different deletion strains (*sir2*Δ, *snf1*Δ, *snf1*Δ *sir2*Δ, *gcn5*Δ, *sir2*Δ *gcn5*Δ, *snf1*Δ *gcn5*Δ) were investigated (unpublished). K83 is located within the malonyl/palmitoyl-CoA transferase (MPT) domain located on the alpha-chain, while the larger proportion of the domain is found on the beta-chain encoded by *FAS1* (Jenni et al., 2007). K173 is located within the acyl carrier protein (ACP) domain of Fas2p, while K1551 is located within the ketoacyl synthase (KS) domain of Fas2p (Lomakin et al., 2007).

FA biosynthesis has been shown to be regulated on multiple levels in yeast. For example, both *ACC1* and *FAS1/2* are subjected to transcriptional regulation by the Ino2p/Ino4p complex and activity of Acc1p and the FAS-complex is repressed by addition of exogenous FAs (Chirala, 1992), possibly due to feedback inhibition by acyl-CoA species (Kamiryo et al., 1976); and as mentioned previously, phosphorylation of Acc1p by Snf1p inhibits enzyme activity (Witters and Watts, 1990; Woods et al., 1994). As FA biosynthesis is highly dependent on acetyl-CoA availability, whose concentration in turn influences the acetylation status of proteins, it is likely that there is a regulatory acetylation mechanism controlling FAS activity also in yeast. Thus, the objective of our study was to investigate the function of the three acetylation sites in Fas2p, confirmed by two independent studies, by exchanging these lysine residues to different AAs. Several studies suggest that lysine residues can be replaced with glutamine (Q), which abolishes the positive charge, or arginine (R), which retains the positive charge, in order to mimic a constitutively acetylated and non-acetylated state, respectively (Megee et al., 1990; Schwer et al., 2006; Wang et al., 2014). By using this residue replacement system, the functional impact of protein acetylation can be studied.

## Methods

### Strains and cultivation conditions

Strains investigated in this study were constructed in the yeast strain background CEN.PK113- 7D and CEN.PK113-5D *faa1*Δ *faa4*Δ (Bergman et al., unpublished). In **Table 1**, the genotype and origin of all strains used and constructed in this study are listed.

**Table 1.** Genotype and origin of strains used and constructed.

| Strain | Genotype | Origin |
| --- | --- | --- |
| CEN.PK113-7D | *MATa MAL2-8c SUC2* | P. Kötter, University of Frankfurt, Germany |
| CEN.PK113-5D ΔΔ | *MATa MAL2-8c SUC2 ura3-52 faa1Δ faa4Δ* | Bergman et al. (unpublished) |
| AB31 | *MATa MAL2-8c SUC2 FAS2<sup>K83Q</sup>* | This study |
| AB32 | *MATa MAL2-8c SUC2 FAS2<sup>K83R</sup>* | This study |
| AB33 | *MATa MAL2-8c SUC2 FAS2<sup>K173Q</sup>* | This study |
| AB34 | *MATa MAL2-8c SUC2 FAS2<sup>K173R</sup>* | This study |
| AB35 | *MATa MAL2-8c SUC2 FAS2<sup>K1551Q</sup>* | This study |
| AB36 | *MATa MAL2-8c SUC2 FAS2<sup>K1551R</sup>* | This study |
| AB37 | *MATa MAL2-8c SUC2 ura3-52 faa1Δ faa4Δ FAS2<sup>K83Q</sup>* | This study |
| AB38 | *MATa MAL2-8c SUC2 ura3-52 faa1Δ faa4Δ FAS2<sup>K83R</sup>* | This study |
| AB39 | *MATa MAL2-8c SUC2 ura3-52 faa1Δ faa4Δ FAS2<sup>K173Q</sup>* | This study |
| AB40 | *MATa MAL2-8c SUC2 ura3-52 faa1Δ faa4Δ FAS2<sup>K173R</sup>* | This study |
| AB41 | *MATa MAL2-8c SUC2 ura3-52 faa1Δ faa4Δ FAS2<sup>K1551Q</sup>* | This study |
| AB42 | *MATa MAL2-8c SUC2 ura3-52 faa1Δ faa4Δ FAS2<sup>K1551R</sup>* | This study |
| AB43 | *MATa MAL2-8c SUC2 ura3-52 faa1Δ faa4Δ FAS2<sup>K83Q, K173Q, K1551Q</sup>* | This study |
| AB44 | *MATa MAL2-8c SUC2 ura3-52 faa1Δ faa4Δ FAS2<sup>K83R, K173R, K1551R</sup>* | This study |

Strains to be transformed were grown in YPD (10 g/L yeast extract, 20 g/L peptone and 20 g/L glucose) before they were harvested and made competent. After transformation, strains were grown on YPD plates containing the antibiotic G418 sulphate at a concentration of 200 mg/L.

Cultivation of yeast strains to determine FA / free fatty acid (FFA) content were conducted in defined minimal medium containing 7.5 g L^−1^ (NH_4_)_2_SO_4_, 14.4 g L^−1^ KH_2_PO_4_ and 0.5 g L^−1^ MgSO_4_. The pH was adjusted to 6.5 with 5 M KOH. Sterile solutions of trace metal and vitamin solutions, as previously described by Verduyn et al. (1992), and sterile carbon source solutions were added after autoclavation. As substrate were used: either 20 or 30 g L^−1^ glucose, either 20 mL L^−1^ or 30 mL L^−1^ ethanol (99,9%), or 10 g L^−1^ glucose in combination with addition of feed beads (12 mm diameter (Adolf Kühner AG) corresponding to approximately 20 g L^−1^ (added after 24 h of cultivation).

For cultivation of strains with origin in CEN.PK113-5D, 60 mg/L uracil was added to the minimal medium.

Cultivations of strains with CEN.PK113-7D background were made in biological duplicates. Strains were pre-cultivated overnight in 15 mL cultivation tubes containing 5 mL of minimal medium, starting from a fresh single colony for each cultivation. The pre-cultures were used to inoculate 50 mL cultures in unbaffled 250 mL shake flasks at an initial OD_600_ of 0.1. Samples for monitoring cell growth were taken throughout the exponential phase to be able to determine μ(max), while samples for FA analysis were taken at three stages: 1) mid-exponential phase, 2) early ethanol phase and 3) stationary phase. For FA analysis, the samples were centrifuged, the cells were washed with PBS, and the cell pellets were freeze dried and stored at −80°C until being used for sample preparation.

Cultivations of strains with CEN.PK113-5D background were made in biological triplicates or quadruplicates. Strains were pre-cultivated overnight in 15 mL cultivation tubes containing 3 mL of minimal medium, starting from a fresh single colony for each cultivation. The pre-cultures were used to inoculate 20 mL cultures in unbaffled 100 mL shake flasks at an initial OD_600_ of 0.1. Samples were taken at two stages: 1) close to glucose depletion and 2) stationary phase (72 h) by withdrawing cells and media and immediately storing the samples at −20°C until used for sample preparation.

### Strain construction

Point mutations (PMs) were introduced into *FAS2* using a plasmid-based CRISPR/Cas9 system. A complete list of plasmids used and constructed can be found in **Table 2**.

**Table 2.** Description and origin of plasmids used and constructed in this study.

| Plasmid | Description | Origin |
| --- | --- | --- |
| pCfB2311 | 2μ origin; P<sub>SNR52</sub>-gRNA-T<sub>SUP4</sub>; natMX-marker | Stovicek et al. (2015) |
| pCfB2312 | CEN/ARS origin; P<sub>TEF1</sub>-cas9-T<sub>CYC1</sub>; kanMX-marker | Stovicek et al. (2015) |
| pgRNA.FAS2.K83 | pCfB2312 P<sub>SNR52</sub>-gRNA(FAS2.K83)-T<sub>SUP4</sub> | This study |
| pgRNA.FAS2.K173 | pCfB2312 P<sub>SNR52</sub>-gRNA(FAS2.K173)-T<sub>SUP4</sub> | This study |
| pgRNA.FAS2.K1551 | pCfB2312 P<sub>SNR52</sub>-gRNA(FAS2.K1551)-T<sub>SUP4</sub> | This study |

The gRNA target sites were selected using the CRISPy tool (Jakočiūnas et al., 2015), as it permits the user to view the entire ORF and where the specific gRNA sequences are located. For all targets, the aim was to select a gRNA which had a low probability of binding off-target, but it had to be a compromise between the requirement of proximity to the site to be point mutated and to the cut site. Furthermore, choice of gRNA sites was also influenced by the option to silently mutate the gRNA/PAM site with few alterations. The chosen gRNA-sites are listed in supplementary **Table S1**.

Primers used to PCR amplify gRNA-cassettes are listed in supplementary **Table S2**. gRNA cassettes were amplified using plasmid CfB2311 (Stovicek et al., 2015), containing a gRNA cassette with the *SNR52* promoter and *SUP4* terminator, as initial template. Primers were designed to amplify the gRNA cassette in two parts, where the 20 bp specific targeting sequence was added to primers as homologous overhang at the reverse primer of the first fragment (containing the *SNR52* promoter) and at the forward primer for the second fragment (containing the structural part of the gRNA and the *SUP4* terminator), generating two DNA products that were fused to a complete gRNA cassette in a second PCR reaction (using primers named F/R- gRNA-HR). PCR reactions were carried out with Phusion high fidelity DNA polymerase (Thermo Fisher Scientific) and PCR products were purified with a Gene Jet PCR purification Kit (Thermo Fisher Scientific).

The centromeric plasmid pCfB2312 (Stovicek et al., 2015) was used in this study to co-express both, Cas9 and guide RNA. The vector was linearized with the restriction enzyme *Pfl*23*II* (Thermo Fisher Scientific) and gRNA-cassettes, flanked during PCR with 35 bp arms homologous to the digested plasmid, were inserted using Gibson assembly^®^ cloning (New England Biolabs). After Gibson assembly^®^ cloning, the mix was used to transform chemically competent *E. coli* cells, according to Inoue et al. (1990). Colonies from each sample were used to inoculate 5 mL LB-medium with ampicillin (80 mg L^−1^) and grown overnight at 37°C, after which plasmids were extracted from the cells using a GeneJET Plasmid Miniprep Kit (Thermo Fisher Scientific). The plasmids were sent for sequencing (Eurofins Genomics), and plasmids with confirmed sequences, named pgRNA.FAS2.K83, pgRNA.FAS2.K173 and pgRNA.FAS2.K1551, were used for downstream modification procedures.

Transformations were carried out using the LiAc/SS carrier DNA/PEG method (Gietz and Schiestl, 2007) with freshly prepared cells. 1 μg of pgRNA-plasmid and 1 nmol of each DS-repair oligo were used in transformations to achieve a PM, and each pgRNA-plasmid was also transformed without DS-oligo as a control. DS-oligos were designed to have their center between Cas9-cut-site and PM-target site and had homology arms of at least 24 bp up- and downstream of the modifications to be introduced. Sequences of repair oligonucleotides can be found in supplementary **Table S3**. Colonies were re-streaked twice on YPD+G418-plates, before the targeted sites were amplified with Phusion high fidelity polymerase and sent for sequencing (Eurofins Genomics) to confirm the PMs. Correct mutants were grown in 5 mL YPD overnight, streaked on YPD plates and incubated at 30°C until single colonies were visible. Afterwards, 10 colonies were replica-plated on YPD and YPD+G418 plates to identify colonies which had lost the gRNA/Cas9 plasmid. In the case of constructing triple glutamine or arginine mutants, three sequential rounds of modifications were carried out.

### Sample preparation and FAME analysis using GC-MS ISQ or GC-FID

In order to quantify intracellular FAs in strains with CEN.PK113-7D background, cell samples were treated according to a previously established protocol by Khoomrung et al. (2012), in which FAs are trans-esterified in methanol to fatty acid methyl esters (FAMEs) (Khoomrung et al., 2012). Approximately 10 mg of freeze dried cell mass and 10 μg of heptadecanoic acid (internal standard, dissolved in hexane) were added to a glass-extraction tube, followed by 1 mL of hexane and 2 mL of 14% BF3 in methanol. The tube was flushed with nitrogen for 30 sec before being capped and vortexed for 20 sec. After this, the remaining microwave assisted FAME-derivatization steps and GC-MS ISQ analysis and quantification procedure followed Khoomrung et al. (2012) precisely.

FFAs produced by strains with CEN.PK113-5D *faa1*Δ*faa4*Δ background were simultaneously extracted and derivatized to FAMEs using a protocol adapted from Haushalter et al. (2014). 300 μL of cell culture (media + cells) were transferred to a glass vial, to which 15 μL of tetrabutylammonium hydroxide were added. Immediately, 300 μL of 200 mM methyl iodide and 100 mg L^−1^ heptadecanoic acid (internal standard) in dichloromethane were added, after which the tube was capped, and samples shaken in a vortex mixer for 30 min at 1400 rpm. The samples were centrifuged to promote phase separation after which 200 μL of the dichloromethane layer (bottom) were transferred to a GC-vial with glass insert, evaporated to dryness, and resuspended in 200 μl hexane. The FAME-samples were analyzed using gas chromatography (Focus GC, ThermoFisher Scientific) equipped with a Zebron ZB-5MS GUARDIAN capillary column (30 m × 0.25 mm × 0.25 μm, Phenomenex) and a Flame Ionization Detector (FID, ThermoFisher Scientific). The inlet temperature was 280°C, and injection volume was 1 μL. The following program was used: initial temperature 50°C, hold for 2 min; ramp to 140°C during 3 min; ramp to 280°C during 14 min; hold for 3 min. The flow rate of helium (carrier gas) was 1.0 mL/min. Six concentrations of FAME-standards (of physiologically relevant length: C12-C18) between 6.25 and 200 mg L^−1^ were analyzed alongside the samples in order to quantify the produced FAs using the Xcalibur software.

### HPLC

In order to verify that the majority of the substrate was consumed in the samples, extracellular metabolites were quantified using high performance liquid chromatography. Fermentation samples were centrifuged and 15 μL of the supernatant were injected into an Ultimate 3000 (Dionex) HPLC system, equipped with an Aminex HPX-87H ion exclusion column (Biorad). The column temperature was set to 45°C, the eluent consisted of 5 mM H_2_SO_4_, and the flow rate was 0.6 mL min^−1^. Glucose, glycerol, acetate and ethanol were detected using a refractive index detector (512 μRIU).

## Results

### Growth behavior and FA levels of CEN.PK113-7D strains point-mutated in FAS2

To investigate the function of three lysine acetylation sites (K83, K173 and K1551) in the Fas2p of *S. cerevisiae*, which have been identified in two independent studies (Henriksen et al., 2012; Kumar et al., unpublished), we modified these sites either to glutamine (Q), to mimic a constitutively acetylated state, or arginine (R) to mimic a non-acetylatable state. This resulted in six CEN.PK113-7D strains, carrying one out of six modified *FAS2* versions, namely *FAS2*^K83Q^, *FAS2*^K83R^, *FAS2*^K173Q^, *FAS2*^K173R^, *FAS2*^K1551Q^ or *FAS2*^K1551R^. Hereafter, the strains are referred to as the AA replacements made, e.g. K83R. These strains and the unmodified parent strain (control) were investigated for their growth behavior as well as their production of FAs when cultivated in minimal medium with 20 g/L of glucose.

The maximal growth rate μ(max) of the strains was between 0.41 and 0.42, whereas the final OD_600_ was between 11.8 and 12.4 (**Table 3**). The μ(max) and the final OD_600_ of the six modified strains were not significantly different (p<0.05) compared to the control strain in which the Fas2p had not been modified.

**Table 3.** Maximum specific growth rate and final OD_600_ of CEN.PK113-7D strains grown in minimal medium containing 2% glucose.

| Strain | $\mu(\text{max})$ [h<sup>-1</sup>]<sup>1</sup> | OD<sub>600</sub>(final)<sup>1</sup> |
| --- | --- | --- |
| Control | 0.419 ± 0.004 | 12.16 ± 0.12 |
| K83Q | 0.416 ± 0.003 | 12.08 ± 0.24 |
| K83R | 0.414 ± 0.008 | 12.04 ± 0.12 |
| K173Q | 0.409 ± 0.005 | 11.78 ± 0.34 |
| K173R | 0.411 ± 0.014 | 12.4 ± |
| K1551Q | 0.409 ± 0.002 | 12.08 ± 0.12 |
| K1551R | 0.418 ± 0.006 | 12.06 ± 0.02 |

<sup>1</sup> The values are means of biological duplicates ± standard deviation (except for the final OD<sub>600</sub> of K173R, which only represents measurement from one replicate).

The total FA content (**Figure 1**) and FA chain composition (**Table 4**) in the control strain as well as the six modified strains were determined in three different growth phases (glucose phase, ethanol phase, stationary phase).

**Figure 1.**
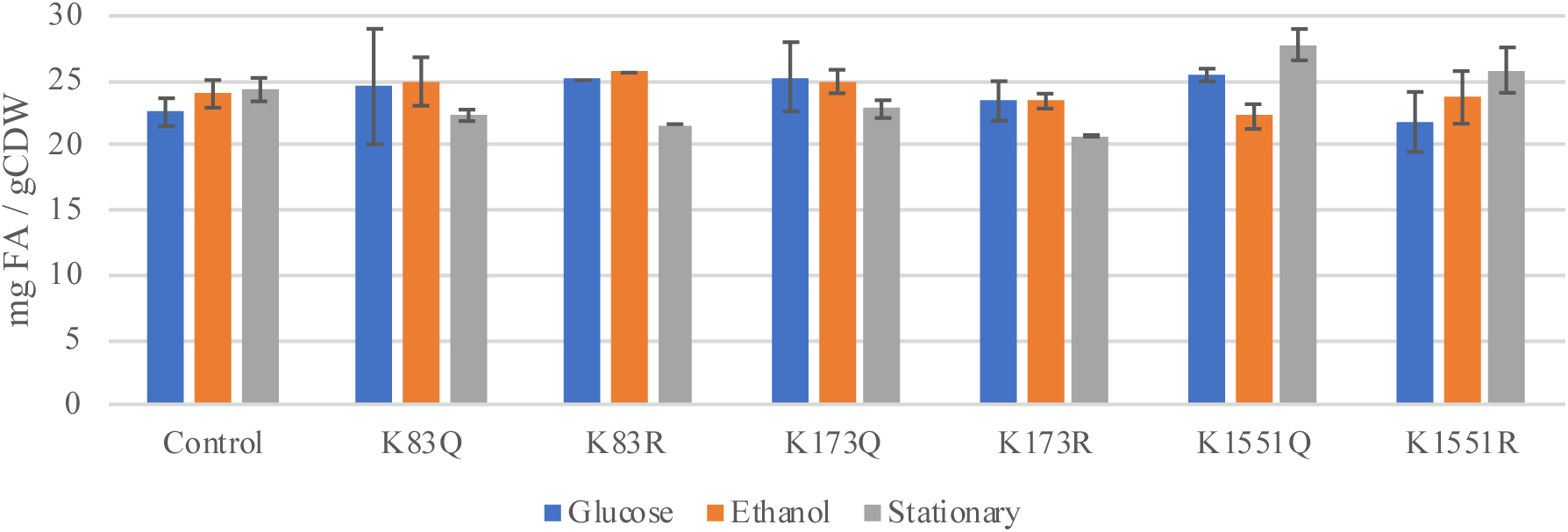
Total fatty acid content (mg/g CDW) in CEN.PK113-7D strains grown in minimal medium, containing 20 g/L glucose harvested in exponential phase (Glucose), early ethanol phase (Ethanol) or stationary phase (Stationary). The values are means of biological duplicates ± standard deviation (except for K83R, which only represents a single measurement, and sample K173R Stationary, which represent duplicate measurements based on biomass from one replicate). gCDW indicates grams of cell dry weight.

**Table 4.**
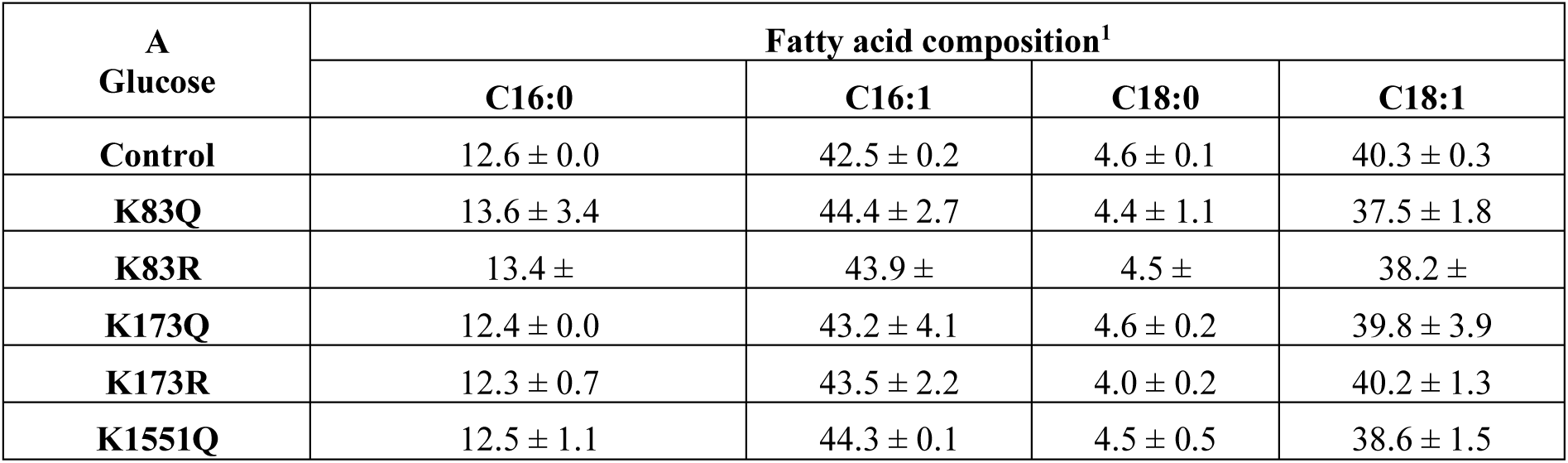
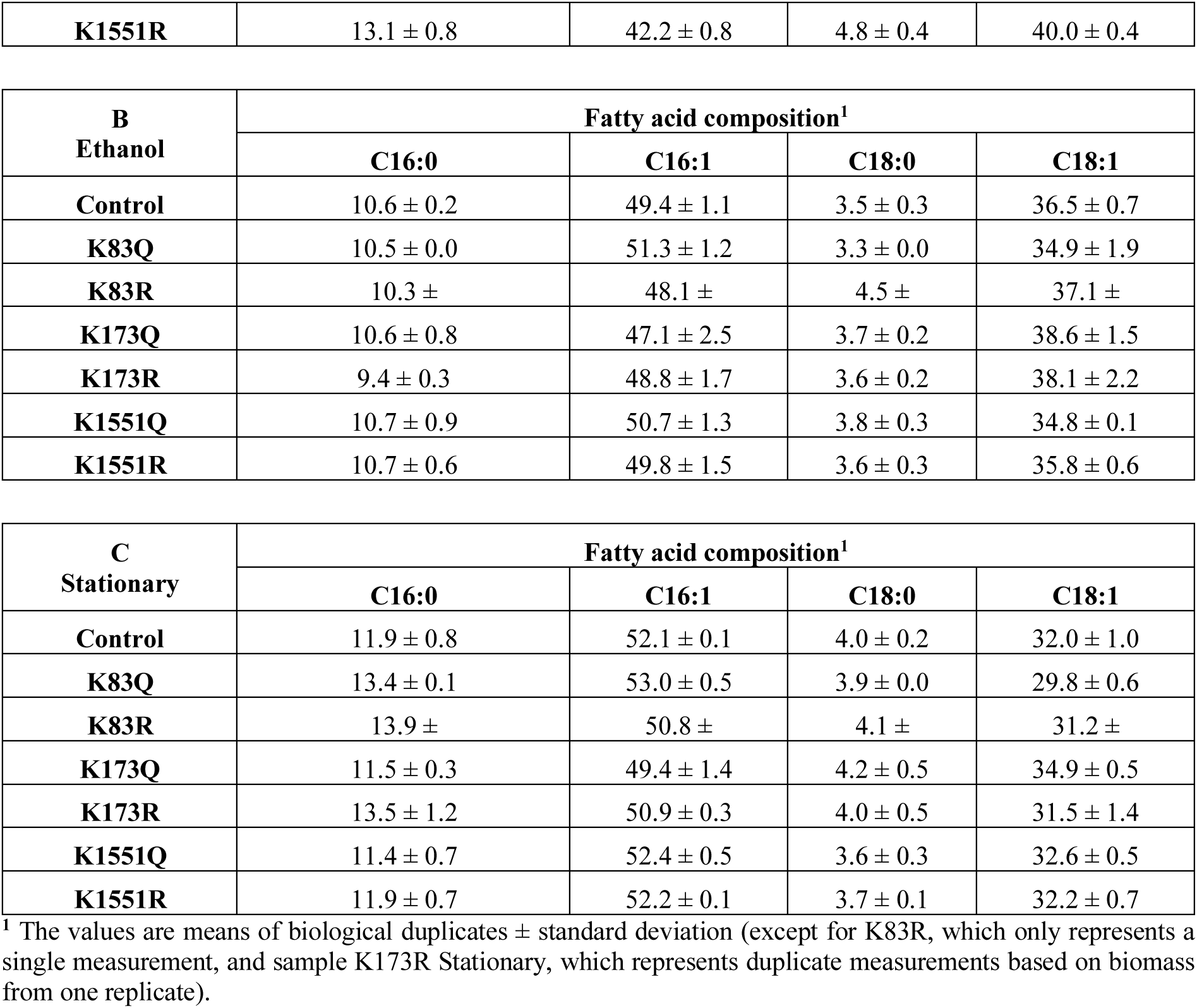
Fatty acid composition (%) of in CEN.PK113-7D strains grown in minimal medium, containing 20 g/L glucose, harvested in exponential phase (A, Glucose), early ethanol phase (B, Ethanol) or stationary phase (C, Stationary).

| A<br>Glucose | Fatty acid composition<sup>1</sup> |  |  |  |
| --- | --- | --- | --- | --- |
|  | C16:0 | C16:1 | C18:0 | C18:1 |
| Control | 12.6 ± 0.0 | 42.5 ± 0.2 | 4.6 ± 0.1 | 40.3 ± 0.3 |
| K83Q | 13.6 ± 3.4 | 44.4 ± 2.7 | 4.4 ± 1.1 | 37.5 ± 1.8 |
| K83R | 13.4 ± | 43.9 ± | 4.5 ± | 38.2 ± |
| K173Q | 12.4 ± 0.0 | 43.2 ± 4.1 | 4.6 ± 0.2 | 39.8 ± 3.9 |
| K173R | 12.3 ± 0.7 | 43.5 ± 2.2 | 4.0 ± 0.2 | 40.2 ± 1.3 |
| K1551Q | 12.5 ± 1.1 | 44.3 ± 0.1 | 4.5 ± 0.5 | 38.6 ± 1.5 |
| **K1551R** | 13.1 ± 0.8 | 42.2 ± 0.8 | 4.8 ± 0.4 | 40.0 ± 0.4 |

| **B**<br>**Ethanol** | **Fatty acid composition<sup>1</sup>** |  |  |  |
| --- | --- | --- | --- | --- |
|  | **C16:0** | **C16:1** | **C18:0** | **C18:1** |
| **Control** | 10.6 ± 0.2 | 49.4 ± 1.1 | 3.5 ± 0.3 | 36.5 ± 0.7 |
| **K83Q** | 10.5 ± 0.0 | 51.3 ± 1.2 | 3.3 ± 0.0 | 34.9 ± 1.9 |
| **K83R** | 10.3 ± | 48.1 ± | 4.5 ± | 37.1 ± |
| **K173Q** | 10.6 ± 0.8 | 47.1 ± 2.5 | 3.7 ± 0.2 | 38.6 ± 1.5 |
| **K173R** | 9.4 ± 0.3 | 48.8 ± 1.7 | 3.6 ± 0.2 | 38.1 ± 2.2 |
| **K1551Q** | 10.7 ± 0.9 | 50.7 ± 1.3 | 3.8 ± 0.3 | 34.8 ± 0.1 |
| **K1551R** | 10.7 ± 0.6 | 49.8 ± 1.5 | 3.6 ± 0.3 | 35.8 ± 0.6 |

| **C**<br>**Stationary** | **Fatty acid composition<sup>1</sup>** |  |  |  |
| --- | --- | --- | --- | --- |
|  | **C16:0** | **C16:1** | **C18:0** | **C18:1** |
| **Control** | 11.9 ± 0.8 | 52.1 ± 0.1 | 4.0 ± 0.2 | 32.0 ± 1.0 |
| **K83Q** | 13.4 ± 0.1 | 53.0 ± 0.5 | 3.9 ± 0.0 | 29.8 ± 0.6 |
| **K83R** | 13.9 ± | 50.8 ± | 4.1 ± | 31.2 ± |
| **K173Q** | 11.5 ± 0.3 | 49.4 ± 1.4 | 4.2 ± 0.5 | 34.9 ± 0.5 |
| **K173R** | 13.5 ± 1.2 | 50.9 ± 0.3 | 4.0 ± 0.5 | 31.5 ± 1.4 |
| **K1551Q** | 11.4 ± 0.7 | 52.4 ± 0.5 | 3.6 ± 0.3 | 32.6 ± 0.5 |
| **K1551R** | 11.9 ± 0.7 | 52.2 ± 0.1 | 3.7 ± 0.1 | 32.2 ± 0.7 |

<sup>1</sup> The values are means of biological duplicates ± standard deviation (except for K83R, which only represents a single measurement, and sample K173R Stationary, which represents duplicate measurements based on biomass from one replicate).

The total FA content in the glucose phase was between 21.8 and 25.4 mg FAs/g cell dry weight (CDW), in the ethanol phase between 22.2 and 25.6 mg FAs/g CDW and in the stationary phase between 20.7 and 27.8 mg FAs/g CDW (**Figure 1**). The FA composition showed that in all strains the most abundant FA was C16:1 (between 42.2% − 53%), followed by C18:1 (between 29.8% − 40.3%), C16:0 (between 9.4% − 13.6%) and C18:0 (between 3.3% − 4.8%) (**Table 4**). Overall, no significant differences (p<0.05) could be observed between the background strain CEN.PK113-7D and the six modified strains for the total FA content or the FA composition. These results indicate that the single AA substitutions in Fas2p at K83, K173 and K1551 in the background strain CEN.PK113-7D do not have an influence on growth behavior, total FA content or FA composition.

### FFA synthesis in CEN.PK113-5D faa1Δ faa4Δ strains point-mutated in FAS2

As the FAS is a highly acetylated protein, we hypothesized that the influence of a single AA residue on enzymatic activity might not be sufficient to determine a regulatory function. Therefore, we also wanted to introduce AA exchanges at all three lysine sites (K83, K173 and K1551) in Fas2p in a single strain, to investigate the effects of the combined mutations. Furthermore, as the product of FA synthesis has multiple cellular destinations (e.g. phospholipids, triacylglycerols, acetyl-CoA via beta-oxidation), we further questioned if it would be feasible to test a regulatory function of acetylation in a strain where a small increase or reduction in FFA production possibly would be masked by other regulatory mechanisms. Since it has been shown that the products of FAS, fatty acyl-CoAs, can inhibit the activity of the enzyme (Chirala, 1992; Kamiryo et al., 1976), we decided to introduce the AA substitutions of Fas2p in the background strain CEN. PK 113-5D *faa1*Δ *faa4*Δ, in which the genes coding for the fatty acyl-CoA synthetases Faa1p and Faa4p are deleted. In such a strain, the inability of FFA activation into acyl-CoA species in part de-regulates FAS-activity, leading to an increased overall flux, FA excretion, and inhibition of consumption of the produced FFA species. This would potentially enhance the effect of a regulatory alteration in Fas2p. Thus, the six single PMs investigated in strain CEN.PK113-7D, namely *FAS2*^K83Q^, *FAS2*^K83R^, *FAS2*^K173Q^, *FAS2*^K173R^, *FAS2*^K1551Q^ or *FAS2*^K1551R^, and the corresponding triple mutants, denoted *FAS2*^QQQ^ and *FAS2*^RRR^, were introduced into CEN.PK113-5D *faa1*Δ *faa4*Δ. These strains were investigated for their total FFA production capacity and FA chain composition during growth on different carbon sources and/or feeding strategies.

Initially, strains were cultivated in biological triplicates using 2% glucose as substrate. Samples were harvested when the strains stopped growing exponentially, indicating that glucose was depleted, and after 72 h, at which the cells had reached stationary phase. In addition, the control strain and the triple mutants were grown in biological triplicates in 2% (v/v) ethanol. The amount of FAs (normalized to OD_600_ for each sample at the time of harvest) produced by the strains grown in glucose are shown in **Figure 2A** and the strains grown in ethanol shown in **Figure 2B**. The composition of the produced FAs at glucose depletion and stationary phase of glucose grown cells are shown in **Table 5A and 5B**, respectively. The composition of the produced FAs in strains grown on ethanol are shown in **Table 5C**.

**Figure 2.**
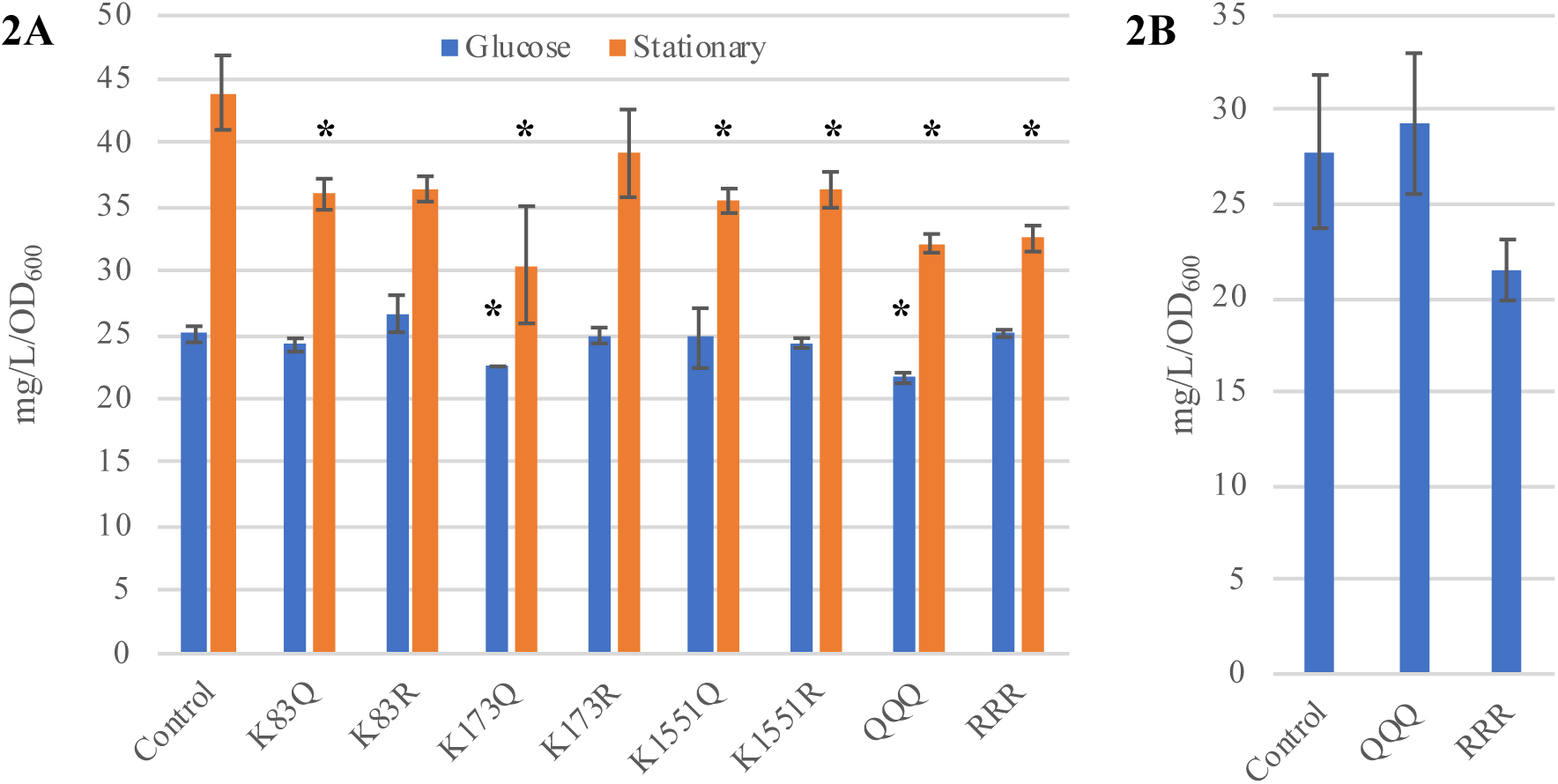
Total free fatty acid content (mg/L/OD_600_) in CEN.PK113-5D *fac1*Δ *faa4*Δ strains grown in minimal medium, **(A)** containing 2% glucose harvested around glucose depletion (Glucose) or after 72 h (Stationary) and **(B)** containing 2% (v/v) ethanol harvested at 72 h. The values are means of biological triplicates ± standard deviation (except for strain K173Q, which represent duplicate measurements). Significant changes (p<0.05. Students t-test, two-tailed, unequal variance assumed) compared to control are marked with an asterisk (*).

**Table 5.** Fatty acid composition (%) in CEN.PK113-5D *faa1*Δ *faa4*Δ strains grown in either minimal medium containing 2% glucose harvested around glucose depletion (**A, Glucose**) or at 72 h (**B, Stationary**), or grown in minimal medium containing 2% (v/v) ethanol harvested at 72 h (**C, Ethanol**).

| **A**<br>**Glucose** | **Fatty acid composition<sup>1</sup>** |  |  |  |  |  |  |  |
| --- | --- | --- | --- | --- | --- | --- | --- | --- |
|  | **C8** | **C10** | **C12** | **C14** | **C16:0** | **C16:1** | **C18:0** | **C18:1** |
| **Control** | 0.3 ± 0.2 | 1.2 ± 0.2 | 0.8 ± 0.1 | 1.2 ± 0.2 | 37.6 ± 0.4 | 26.8 ± 0.2 | 23.3 ± 0.9 | 8.8 ± 0.1 |
| **K83Q** | 0.6 ± 0.6 | 0.7 ± 0.3 | 0.6 ± 0.1 | 0.7 ± 0.3 | 38.0 ± 1.1 | 26.7 ± 0.4 | 22.5 ± 1.6 | 10.1 ± 0.4 |
| **K83R** | 0.8 ± 0.4 | 0.9 ± 0.1 | 0.7 ± 0.0 | 0.9 ± 0.1 | 35.6 ± 1.2 | 26.4 ± 0.7 | 24.8 ± 2.0 | 9.9 ± 0.3 |
| **K173Q** | 1.6 ± 0.3 | 1.4 ± 0.0 | 0.8 ± 0.1 | 1.4 ± 0.0 | 37.6 ± 1.2 | 19.0 ± 4.0 | 29.1 ± 2.9 | 9.2 ± 0.4 |
| **K173R** | 1.1 ± 0.7 | 1.1 ± 0.2 | 0.8 ± 0.0 | 1.1 ± 0.2 | 38.2 ± 1.2 | 25.1 ± 1.1 | 23.1 ± 1.2 | 9.4 ± 0.7 |
| **K1551Q** | 0.2 ± 0.2 | 0.8 ± 0.2 | 0.6 ± 0.0 | 0.8 ± 0.2 | 37.3 ± 1.3 | 25.0 ± 1.9 | 26.4 ± 1.8 | 8.8 ± 0.8 |
| **K1551R** | 0.8 ± 0.4 | 0.9 ± 0.1 | 0.7 ± 0.0 | 0.9 ± 0.1 | 38.4 ± 1.2 | 26.6 ± 0.5 | 21.7 ± 0.9 | 9.9 ± 0.1 |
| **QQQ** | 0.8 ± 0.5 | 1.1 ± 0.2 | 0.7 ± 0.1 | 1.1 ± 0.2 | 36.5 ± 1.6 | **22.3 ± 0.5** | **27.8 ± 1.4** | 9.6 ± 0.5 |
| **RRR** | 0.8 ± 0.7 | 1.3 ± 0.2 | 0.9 ± 0.1 | 1.3 ± 0.2 | 36.9 ± 0.5 | 26.6 ± 0.7 | 22.9 ± 1.5 | 9.4 ± 0.4 |

| **B**<br>**Stationary** | **Fatty acid composition<sup>1</sup>** |  |  |  |  |  |  |  |
| --- | --- | --- | --- | --- | --- | --- | --- | --- |
|  | **C8** | **C10** | **C12** | **C14** | **C16:0** | **C16:1** | **C18:0** | **C18:1** |
| **Control** | 2.1 ± 0.1 | 0.0 ± 0.0 | 0.0 ± 0.0 | 0.0 ± 0.0 | 29.2 ± 0.7 | 23.5 ± 1.0 | 34.3 ± 0.4 | 11.0 ± 0.6 |
| **K83Q** | **2.8 ± 0.2** | 0.0 ± 0.0 | 0.0 ± 0.0 | 0.0 ± 0.0 | 29.2 ± 0.4 | 23.2 ± 0.5 | 33.5 ± 0.8 | 11.2 ± 0.3 |
| **K83R** | 2.7 ± 0.5 | 0.0 ± 0.0 | 0.0 ± 0.0 | 0.0 ± 0.0 | **25.9 ± 1.1** | 24.2 ± 1.8 | 35.2 ± 0.3 | 12.1 ± 0.2 |
| **K173Q** | 3.6 ± 1.1 | 0.0 ± 0.0 | 0.0 ± 0.0 | 0.0 ± 0.0 | **21.9 ± 0.6** | 16.8 ± 4.0 | 45.1 ± 4.1 | 12.6 ± 0.5 |
| **K173R** | 2.3 ± 0.3 | 0.0 ± 0.0 | 0.0 ± 0.0 | 0.0 ± 0.0 | **26.5 ± 1.0** | 22.7 ± 1.5 | 36.2 ± 1.0 | 12.3 ± 0.5 |
| **K1551Q** | **2.7 ± 0.1** | 0.0 ± 0.0 | 0.0 ± 0.0 | 0.0 ± 0.0 | **24.9 ± 1.2** | 24.7 ± 1.3 | **35.9 ± 0.1** | 11.9 ± 0.1 |
| **K1551R** | **2.5 ± 0.1** | 0.0 ± 0.0 | 0.0 ± 0.0 | 0.0 ± 0.0 | **26.6 ± 0.1** | 22.9 ± 1.1 | 36.1 ± 1.1 | 12.0 ± 0.0 |
| **QQQ** | 1.7 ± 0.4 | 0.0 ± 0.0 | 0.0 ± 0.0 | 0.0 ± 0.0 | **25.2 ± 0.6** | 22.2 ± 0.2 | **38.5 ± 0.8** | 12.3 ± 0.1 |
| **RRR** | 1.8 ± 0.3 | 0.0 ± 0.0 | 0.0 ± 0.0 | 0.0 ± 0.0 | **26.3 ± 1.0** | 21.2 ± 1.1 | **37.8 ± 0.2** | **12.9 ± 0.2** |

| **C**<br>**Ethanol** | **Fatty acid composition<sup>1</sup>** |  |  |  |  |  |  |  |
| --- | --- | --- | --- | --- | --- | --- | --- | --- |
|  | **C8** | **C10** | **C12** | **C14** | **C16:0** | **C16:1** | **C18:0** | **C18:1** |
| **Control** | 1.9 ± 0.2 | 0.0 ± 0.0 | 0.0 ± 0.0 | 0.0 ± 0.0 | 20.4 ± 2 | 31.8 ± 3.7 | 31.9 ± 1.8 | 14 ± 0.3 |
| **QQQ** | 1.6 ± 1.1 | 0.0 ± 0.0 | 0.0 ± 0.0 | 0.0 ± 0.0 | 20.3 ± 3.8 | 30.8 ± 4.9 | 31.8 ± 3 | 15.5 ± 1.6 |
| **RRR** | 1.1 ± 1.0 | 0.0 ± 0.0 | 0.0 ± 0.0 | 0.0 ± 0.0 | 18.4 ± 1.6 | 32.9 ± 2.7 | 32.7 ± 2.3 | 15 ± 0.1 |

<sup>1</sup> The values are means of biological triplicates ± standard deviation (except for strain K173Q, which represent duplicate measurements). Significant changes (p<0.05, Students t-test, two-tailed, unequal variance assumed) compared to control are marked in bold.

Based on strains grown in glucose media and samples harvested around glucose depletion (Glucose, **Figure 2A**), lysine to glutamine modification resulted in a reduction of FFA levels (mg/L/OD_600_) in strains K173Q and QQQ, corresponding to 90% and 86% of the control, while corresponding alterations to arginine did not result in a significant decrease in FFA concentrations. When considering samples harvested at stationary phase (Stationary, **Figure 2A**), all strains with *FAS2*-PMs except for K83R and K173R showed a significant reduction in FFA level, where the strongest reduction was seen for K173Q, QQQ and RRR – 69%, 73% and 74%) of the level seen for the control. The triple mutation strains (QQQ/RRR) did not display any significant changes in FFA concentrations compared to the control when growing on ethanol (**Figure 2B**). Strain RRR did show a moderate reduction compared to the control (p-value = 0.14), however it should be noted that the error bars for the control strain were large.

With regard to FA composition for strains grown in glucose, the only strain resulting in a significantly skewed FA composition around glucose depletion was the triple glutamine mutant QQQ, which had a reduced amount of C16:1 and increased amount of C18:0 (**Table 5A**). The same trends can be observed for strain K173Q, but at a non-significant level. In the stationary phase, several Fas2p mutants showed a reduction in C16:0 (all strains but K83Q), and K1551Q, QQQ and RRR showed a significant increase in C18:0 species (**Table 5B**). Strain RRR also had a significant increase in C18:1 FAs. The triple mutants grown in ethanol did not show a significant difference in FA distribution compared to the control (**Table 5C**).

In order to obtain more reliable results, we decided to repeat the experiment using 3% glucose or 3% (v/v) ethanol as substrate to increase production which potentially could facilitate seeing differences imposed by the Fas2p-modifications, and conduct the experiment using biological quadruplicates. Furthermore, we wanted to investigate if the modifications would result in a difference when strains were grown in glucose-limited conditions using so-called feed beads (corresponding to 2% glucose) after a shorter initial growth period (using 1% dissolved glucose). The amount of FAs (normalized to OD_600_ for each sample at the time of harvest) produced by the strains grown in glucose are shown in **Figure 3A** and the strains grown in ethanol and using feed beads are shown in **Figure 3B**. The composition of the produced FFAs at glucose depletion and stationary phase of glucose grown cells are shown in **Table 6A and 6B**, respectively. The composition of the produced FFAs in strains grown on ethanol and using feed beads are shown in **Table 6C and 6D**, respectively.

**Figure 3.**
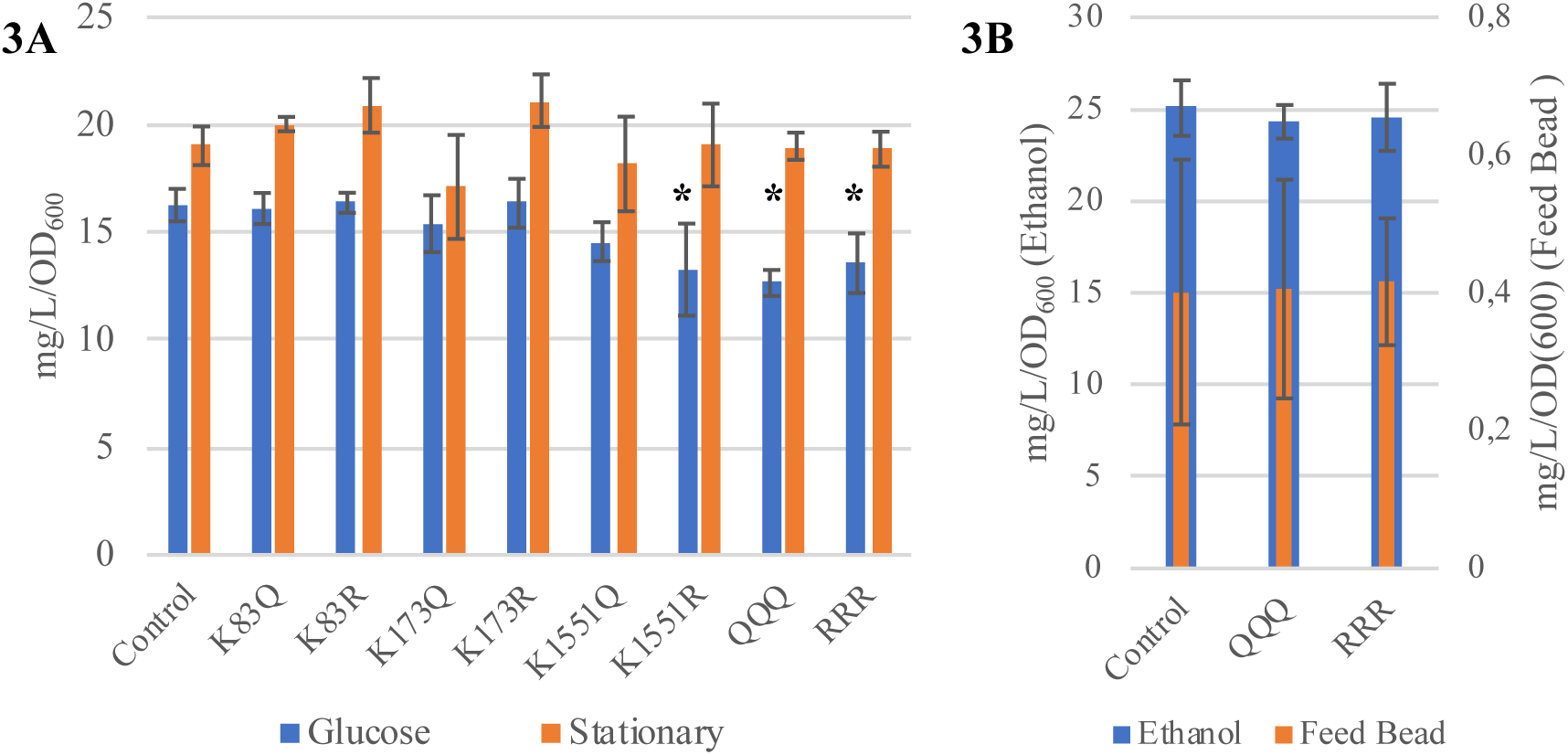
Total free fatty acid content (mg/L/OD_600_) in CEN.PK113-5D *fact1*Δ *faa4*Δ strains grown in minimal medium, **(A)** containing 3% glucose harvested around glucose depletion (Glucose) or after 72 h (Stationary) and **(B)** containing 3% (v/v) ethanol harvested at 72 h (Ethanol) or 1% dissolved glucose and 2% glucose supplied in feed bead format added after 24 h of culture, harvested after 96 h (Feed Bead). The values are means of biological quadruplicates ± standard deviation (except for K173R, which represents triplicate measurements). Significant changes (p<0.05, Students t-test, two-tailed, unequal variance assumed) compared to control marked with an asterisk (*).

**Table 6.** Fatty acid composition (%) in CEN.PK113-5D *faa1*Δ *faa4*Δ strains grown in either minimal medium containing 3% glucose harvested around glucose depletion (**A, Glucose**) or at 72 h (**B, Stationary**), in minimal medium containing 3% (v/v) ethanol harvested at 72 h (**C, Ethanol**), or minimal medium containing 1% dissolved glucose and 2% glucose supplied in feed bead format added after 24 h of culture, harvested after 96 h (**D, Feed Bead**).

| A<br>Glucose | Fatty acid composition<sup>1</sup> |  |  |  |  |  |  |  |
| --- | --- | --- | --- | --- | --- | --- | --- | --- |
|  | C8 | C10 | C12 | C14 | C16:0 | C16:1 | C18:0 | C18:1 |
| Control | 0.0 ± 0.0 | 0.5 ± 0.1 | 0.5 ± 0.1 | 0.4 ± 0.1 | 27.6 ± 3.3 | 18.9 ± 0.8 | 40.0 ± 2.8 | 12.3 ± 0.5 |
| K83Q | 0.1 ± 0.1 | 0.5 ± 0.0 | 0.5 ± 0.0 | 0.3 ± 0.0 | 27.1 ± 1.2 | 19.6 ± 1.5 | 39.3 ± 3.3 | 12.6 ± 0.9 |
| K83R | 0.0 ± 0.0 | 0.5 ± 0.0 | 0.5 ± 0.0 | 0.4 ± 0.0 | 27.2 ± 1.1 | 19.2 ± 0.4 | 40.7 ± 1.1 | 11.6 ± 0.3 |
| K173Q | 1.1 ± 1.5 | **0.7 ± 0.1** | **0.7 ± 0.1** | 0.4 ± 0.1 | 24.9 ± 2.2 | **13.0 ± 1.8** | **48.0 ± 2.9** | 11.2 ± 0.7 |
| K173R | 0.1 ± 0.1 | 0.6 ± 0.1 | 0.6 ± 0.1 | 0.4 ± 0.0 | 26.8 ± 1.4 | 19.1 ± 0.5 | 40.3 ± 1.1 | 12.1 ± 0.4 |
| K1551Q | 0.2 ± 0.2 | 0.5 ± 0.1 | 0.5 ± 0.1 | 0.3 ± 0.1 | 26.9 ± 4.0 | 18.0 ± 0.9 | 41.1 ± 3.8 | 12.5 ± 0.5 |
| K1551R | 0.1 ± 0.1 | 0.6 ± 0.1 | 0.6 ± 0.1 | 0.4 ± 0.1 | 27.7 ± 2.5 | 20.0 ± 0.9 | 37.4 ± 3.7 | 13.3 ± 1.2 |
| QQQ | 0.1 ± 0.1 | 0.4 ± 0.1 | 0.4 ± 0.1 | 0.3 ± 0.0 | 29.5 ± 2.1 | **14.5 ± 0.9** | 43.8 ± 1.5 | **11.0 ± 0.5** |
| RRR | 0.4 ± 0.4 | 0.7 ± 0.2 | 0.7 ± 0.2 | **0.6 ± 0.1** | 30.5 ± 0.8 | 18.0 ± 0.5 | 37.4 ± 0.8 | 11.7 ± 0.5 |

| B<br>Stationary | Fatty acid composition<sup>1</sup> |  |  |  |  |  |  |  |
| --- | --- | --- | --- | --- | --- | --- | --- | --- |
|  | C8 | C10 | C12 | C14 | C16:0 | C16:1 | C18:0 | C18:1 |
| Control | 1.7 ± 0.1 | 0.1 ± 0.0 | 0.1 ± 0.0 | 0.0 ± 0.0 | 28.5 ± 0.4 | 24.2 ± 0.6 | 35.5 ± 0.5 | 9.9 ± 0.4 |
| K83Q | 1.8 ± 0.3 | **0.0 ± 0.0** | **0.0 ± 0.0** | 0.0 ± 0.0 | **30.1 ± 0.4** | **26.5 ± 0.1** | **31.5 ± 0.3** | 10.2 ± 0.2 |
| K83R | 1.9 ± 0.7 | **0.0 ± 0.0** | **0.0 ± 0.0** | 0.0 ± 0.0 | 29.8 ± 1.6 | 25.4 ± 1.0 | **32.5 ± 0.6** | 10.4 ± 0.4 |
| K173Q | 2.6 ± 0.8 | **0.0 ± 0.0** | **0.0 ± 0.0** | 0.0 ± 0.0 | 28.3 ± 0.8 | **18.0 ± 2.2** | **40.9 ± 1.6** | 10.1 ± 0.8 |
| K173R | 2.0 ± 0.3 | **0.0 ± 0.0** | **0.0 ± 0.0** | 0.0 ± 0.0 | 28.2 ± 0.4 | 24.7 ± 0.3 | **34.4 ± 0.4** | **10.7 ± 0.2** |
| K1551Q | 1.9 ± 0.3 | **0.0 ± 0.0** | **0.0 ± 0.0** | 0.0 ± 0.0 | 30.4 ± 1.8 | 24.8 ± 0.9 | **32.6 ± 0.7** | 10.3 ± 0.3 |
| K1551R | 2.3 ± 0.5 | **0.0 ± 0.0** | **0.0 ± 0.0** | 0.0 ± 0.0 | 29.9 ± 1.2 | 24.4 ± 1.2 | **32.8 ± 0.5** | 10.6 ± 0.3 |
| QQQ | **2.2 ± 0.2** | **0.0 ± 0.0** | **0.0 ± 0.0** | 0.0 ± 0.0 | 29.4 ± 0.4 | 23.2 ± 0.6 | 35.3 ± 0.8 | 9.9 ± 0.1 |
| RRR | **2.0 ± 0.1** | **0.0 ± 0.0** | **0.0 ± 0.0** | 0.0 ± 0.0 | **30.6 ± 1.0** | 23.5 ± 1.2 | 34.0 ± 1.1 | 9.9 ± 0.5 |

| C<br>Ethanol | Fatty acid composition<sup>1</sup> |  |  |  |  |  |  |  |
| --- | --- | --- | --- | --- | --- | --- | --- | --- |
|  | C8 | C10 | C12 | C14 | C16:0 | C16:1 | C18:0 | C18:1 |
| Control | 0.9 ± 0.4 | 0.0 ± 0.0 | 0.0 ± 0.0 | 0.0 ± 0.0 | 25.4 ± 1.3 | 37.5 ± 1.3 | 23.7 ± 2.3 | 12.4 ± 0.1 |
| QQQ | 1.2 ± 0.1 | 0.0 ± 0.0 | 0.0 ± 0.0 | 0.0 ± 0.0 | 26.4 ± 1.7 | 33.7 ± 2.4 | 26.5 ± 0.8 | **12.0 ± 0.2** |
| RRR | 1.1 ± 0.4 | 0.0 ± 0.0 | 0.0 ± 0.0 | 0.0 ± 0.0 | 26.0 ± 1.3 | 34.1 ± 4.0 | 26.2 ± 2.3 | 12.5 ± 0.3 |

| D<br>Feed Bead | Fatty acid composition<sup>1</sup> |  |  |  |  |  |  |  |
| --- | --- | --- | --- | --- | --- | --- | --- | --- |
|  | C8 | C10 | C12 | C14 | C16:0 | C16:1 | C18:0 | C18:1 |
| Control | 46.9 ± 11.7 | 0.0 ± 0.0 | 0.0 ± 0.0 | 0.0 ± 0.0 | 22.3 ± 7.3 | 15.7 ± 5.9 | 4.0 ± 1.9 | 11.1 ± 2.4 |
| QQQ | 62.5 ± 7.6 | 0.0 ± 0.0 | 0.0 ± 0.0 | 0.0 ± 0.0 | 14.4 ± 2.2 | 11.0 ± 2.6 | 1.7 ± 1.9 | 10.4 ± 4.1 |
| RRR | 57.2 ± 9.4 | 0.0 ± 0.0 | 0.0 ± 0.0 | 0.0 ± 0.0 | 15.7 ± 4.9 | 10.1 ± 2.8 | 1.9 ± 1.9 | 15.1 ± 4.5 |

<sup>1</sup> The values are means of biological quadruplicates ± standard deviation (except for K173R, which represents triplicate measurements). Significant changes (p<0.05, Students t-test, two-tailed, unequal variance assumed) compared to control are marked in bold.

When strains were grown in glucose media and samples harvested around glucose depletion (Glucose, **Figure 3A**), FFA concentrations were significantly reduced in strains K1551R, QQQ and RRR (82%, 78% and 83% of control), while the results from the previous run (Glucose, **Figure 2A**) showed K173Q and QQQ to have reduced FFA concentrations in glucose phase. When the strains reached stationary phase (Stationary, **Figure 3A**), no significant changes in FFA levels could be detected, which also is contradictory to the results of the first run (Stationary, **Figure 2A**), where most modifications led to reduced FFA concentrations compared to the control. When considering samples grown in either ethanol or using feed beads (Ethanol and Feed Bead, **Figure 3B**), no significant differences in FFA concentrations could be seen. In general, FFA concentrations were lower when a higher concentration of glucose was used (compare **Figure 2A and 3A**). Interestingly, growth under glucose-limitation (**Figure 3B**) drastically decreased FFA production, as FFA concentrations were as low as 0.4 mg/L/OD_600_ compared to 15-25 mg/L/OD_600_ when glucose or ethanol was used as substrate.

The analysis of the FA composition for strains grown in glucose show that strain K173Q and QQQ significantly decreased the pool of C16:1 and that K173Q increased the amount of C18:0 around glucose depletion (**Table 6A**). These results are relatively consistent with the result from the first run (**Table 5A**), where QQQ brought about the same decrease in C16:1 at a significant level. As mentioned, strain K173Q also showed the same trends during the first run, however, as only duplicate measurements could be obtained, the standard deviations were large, likely preventing a significant difference to be observed. At stationary phase, strain K173Q still showed an increased percentage of C18:0 FAs at the expense of C16:1 compared to the control (**Table 6B**). This is notable, since all the other single mutants led to a significantly reduced amount of C18:0. The general reduction of C16:0 species in the Fas2p mutants seen during the first run was not present in the second run, where two strains, K83Q and RRR, on the contrary increased the proportion of C16:0. When grown in ethanol and in glucose-limited conditions, no significant differences in FA composition could be detected (**Table 6C** and **6D**, respectively).

## Discussion

FAS in yeast is a macromolecular assembly with six copies of each Fas1p and Fas2p, which by themselves are large proteins (2051 and 1887 AAs, respectively). These proteins contain the highest amount of unique detected acetylation sites in yeast, and due to their metabolic proximity to acetyl-CoA – a critical determinant of protein acetylation status and the substrate for FA synthesis – it is likely that FAS-activity would be regulated by acetylation.

The three lysine sites investigated in this study were selected based on two separate acetylation data sets. In the study conducted by Henriksen et al. (2012) samples were collected in exponential phase, where acetyl-CoA levels are high (Cai et al., 2011), and in total detected approximately 4000 unique sites of lysine acetylation. Kumar et al. (unpublished) collected their samples under glucose-limited conditions and detected 260 unique acetyl-lysines. Thus, the number of acetyl-sites detected in Fas1p and Fas2p by Henriksen et al. (in total 79) was considerably higher than observed by Kumar et al. (unpublished), where no sites were detected in Fas1p and only the three lysine sites investigated in this study were found for Fas2p. As indicated by Henriksen et al. (2012), the enrichment of acetyl-lysine containing peptides with acetyl-lysine antibodies is very sensitive to batch-variation in terms of efficiency and specificity, which consequently may have partly influenced the enrichment in the study by Kumar et al. (unpublished). Furthermore, it is likely that acetyl-CoA levels are lower in glucose-limited conditions compared to exponential phase, which likely influence the overall degree of protein acetylation. The three sites, which were detected for Fas2p in the data of Kumar et al. (unpublished) could thus potentially represent more specifically acetylated residues, as at least K1551 and K173 were detected in all strains investigated (Supplementary **Table S4**). K83, K173 and K1551 are all located within functional domains of Fas2p. K83 is located within the alpha-chain specific part of the MPT domain (Jenni et al., 2007), K173 is located within the ACP domain and close to the site of attachment of the pantetheine arm (S180), and K1551 is located relatively close to the catalytic triad of the KS domain (C1305, H1542 and H1583) (Lomakin et al., 2007). Furthermore, all three sites are evolutionary conserved among fungal yeast species (Jenni et al., 2007). Taken together, these considerations suggested that K83, K173 and K1551 of Fas2p were good candidates for an initial investigation.

In order to investigate if and how yeast FAS activity is regulated, we used a residue replacement system where the lysine (K) sites previously detected to be acetylated were changed to either glutamine (Q) or arginine (R), a strategy which has been used in previous studies (Megee et al., 1990; Schwer et al., 2006; Wang et al., 2014). Lysine carries a side chain of four aliphatic carbons and an amino group which is positively charged under biological conditions. If acetylated, the acetyl group is added to the amino group, thus abolishing the positive charge. Arginine carries a side chain of three aliphatic carbons and a guanidine group which also is positively charged under biological conditions, thus mimicking a constitutively non-acetylated lysine. Glutamine, one the other hand, carries only two aliphatic carbons on its side chain, followed by an uncharged amide group, mimicking a constitutively acetylated lysine. The use of this replacement system has limitations, as it assumes that lysine functionality in a protein strictly is dependent on charge. Arginine is bulkier than the unmodified lysine residue, while glutamine is smaller than the acetylated lysine, as described above. Thus, the introduction of either of the residues into the three-dimensional protein structure can interfere with catalytic function in an acetylation-independent manner. In order to use the K➔Q/R replacement system to evaluate regulatory impact of acetylation on protein function, preferably, only the K➔Q replacement should generate a difference in enzyme activity while K➔R should remain unchanged compared to the control. This assumes that the natural state of the residue is unacetylated, otherwise the reverse situation could be observed. It should also be highlighted that this only provides a first indication if a lysine residue is involved in regulation of enzyme activity via acetylation, and further studies are required to verify its true impact.

Our initial attempt to introduce single points mutations of K83, K173 and K1551 of Fas2p into the strain CEN.PK113-7D did not reveal any significant changes related to growth nor FA synthesis or FA composition (**Table 3; Figure 1; Table 4**). Fas1p and Fas2p act as a complex and together have been shown to carry 79 unique sites of acetylation under high acetyl-CoA conditions (29 in Fas1p and 50 in Fas2p) (Henriksen et al., 2012). Therefore, it is not unlikely that a regulatory effect of Fas1p/2p-acetylation is dependent on the simultaneous acetylation of many lysine residues. Thus, we wanted to investigate which effect either QQQ or RRR triple mutations of Fas2p would have on FFA concentrations. Furthermore, we speculated that it would be more suitable to study FA synthesis in a genetic background where the FAS is deregulated. Indeed, when the single and triple AA substitutions were introduced into the parent strain CEN.PK113-5D *faa1*Δ *faa4*Δ, where acyl-CoA inhibition is reduced, significant alterations in FFA concentrations and chain length distribution were observed.

The initial FFA quantification results of strain CEN.PK113-5D *faa1*Δ *faa4*Δ at glucose depletion (Glucose, **Figure 2A**) showed that strain QQQ exhibited a significant reduction in FFA titers, while strain RRR did not, which could be an indication that acetylation has a negative effect on FAS-activity when glucose is being consumed. However, as both triple glutamine (QQQ) and triple arginine (RRR) mutants induce a similar reduction in FFA concentrations when sampled in stationary phase (Stationary, **Figure 2A**) and at glucose depletion in the second experiment (Glucose, **Figure 3A**), it is highly likely that both residue replacements interfere with Fas2p catalytic activity. Furthermore, the differences observed in FFA concentrations in the first and second experiment (**Figure 2A and 3A**), particularly at stationary phase, are contradictory, which challenges the reproducibility of the results.

An observed difference which was consistent between the first and the second experiment with strain CEN.PK113.5D *faa1*Δ *faa4*Δ (though, only significant in the second experiment) was that specifically K173Q, but not K173R appeared to influence the chain length distribution relative to the control. During the second experiment, at glucose depletion, C16:1 species were reduced to 13.0% ± 1.8% in K173Q compared to 18.9% ± 0.8% observed for the control, while C18:0 were increased to 48.0% ± 2.9% compared to 40.0% ± 2.8% observed for the control (**Table 6A**). The corresponding relative amounts of C16:0 at stationary phase were 18.0% ± 2.2% in K173Q compared to 24.2% ± 0.6% observed for the control, while C18:0 were increased to 40.9% ± 1.6% compared to 35.5% ± 0,5% observed for the control (**Table 6B**). The same trends were seen for the triple mutant QQQ in both experiments at glucose depletion, but only significant for both species in the first experiment (**Table 5A**) and for C16:0 reduction in the second experiment (**Table 6A**). The reduction in FFA concentrations in the triple mutant is likely to be traced back to the effect of the K173Q modification in this strain, as the other PMs does not influence the FA distribution in this manner.

K173 is located within the ACP domain of FAS, which is instrumental in the FA synthesis process as it carries the growing acyl-chain and transfers it to the different catalytic centers in the FAS-complex (Jenni et al., 2007). Potentially, the introduction of a constitutively neutral charge or a smaller AA at site K173 generates more space for the growing acyl chain within the FAS-chamber, thus increasing the amount of C18 FAs at the expense of C16 FAs. However, a recent study presented that it was possible to add a complete thioesterase domain at the C-terminal end of the *S. cerevisiae* FAS next to the ACP domain without causing loss of function (Zhu et al., 2017). In order to determine if the observed difference in FA chain-length is the result of a regulatory acetylation on K173 or a steric alteration of the catalytic chamber when introducing a non-charged/smaller AA, further studies are required.

The decision to evaluate impact of three lysine sites in a protein reported to be able to carry at least 50 acetylated lysines, and closer to 80 when Fas1p also is taken into consideration, could possibly have prevented the identification true regulatory lysine sites. The K➔Q/R residue-replacement system is of greater value when the investigated protein has a more limited number of detected acetylation-sites. This was demonstrated in the study by Wang et al. (2014), which identified regulatory acetylation sites in human glucose-6-phosphate dehydrogenase by mutating all seven previously detected acetylation sites in the protein (Choudhary et al., 2009), pinpointing one site to be of regulatory type.

An alternative strategy of investigating the impact of regulation is to focus on the general impact of the acetylation on enzymatic activity. The functional FAS is a large complex consisting of α_6_β_6_ heterododecamers, with six catalytic chambers in which each eight enzymatic reactions or transfers are catalyzed (Lomakin et al., 2007). His-tagging of the C-terminus of Fas1p has been reported to not influence enzyme function, and permits for a column-purification of the complete functional FAS-complex (Johansson et al., 2008). By treating growing cells with the compound nicotinamide (NAM), which is an inhibitor of the sirtuin family of deacetylases (Avalos et al., 2005), and/or trichostatin A (TSA), an inhibitor of deacetylases of group I and II (Vigushin et al., 2001), proteins can be purified with an increased level of acetylations. By blotting the purified enzyme from non-treated and NAM/TSA-treated cells with a pan anti-acetyl lysine antibody, one could determine which group of deacetylase enzymes (I/II or III) is responsible for the removal of the acetyl-groups. Furthermore, by setting up an *in vitro* assay for FAS-activity where acetyl-CoA, malonyl-CoA and NADPH are supplied and consumption of NADPH is monitored, the functional impact of general FAS-acetylation status could be resolved. Determining the influence of individual acetylations would however not be possible using this strategy. Such knowledge would be useful for applied purposes, e.g. in order to abolish regulation via substitution of lysine-sites which has particularly negative impact on enzymatic activity in an acetylated state.

By utilizing amber suppression technology, unnatural amino acids like acetyl-lysine can be directly introduced into proteins by using a to the host orthogonal pair of aminoacyl-tRNA synthetase and tRNA (Neumann et al., 2018). In *S. cerevisiae*, this has previously been shown by Hancock et al, which created a functional expression system for a pyrrolysyl-tRNA synthetase/tRNA^Pyl^_CUA_ pair from *Methanosarcina barkeri* and *M. mazei* which was able to incorporate of a range of unnatural amino acids into proteins, including acetyl-lysine (Hancock et al., 2010). Thus, by coexpressing the mentioned heterologous and orthogonal aminoacyl-tRNA synthetase and tRNA_CUA_ in yeast, and simultaneously introducing GAT-codons at sites to be investigated, particular acetyl-lysines can be obtained. However, in contrast to when a glutamine residue is introduced, the yeast will be able to deacetylate an incorporated acetyl-lysine.

As our study was not able to generate data to determine which type of regulatory function the many acetylation sites have on *S. cerevisiae* FAS-function, we can merely speculate if the regulation is of inhibitory or stimulatory type. Based on the two initial acetylation data sets, there is limited information available, as no quantification of FAs has been conducted. The acetylome data of Kumar et al (unpublished), covers six strains in which deletions of the genes encoding the global regulators Snf1, Sir2 and Gcn5 or combinations thereof were made. The degree of acetylation of Fas2p was found to be negative in *sir2*Δ and *snf1*Δ strains, while it was positive in *gcn5*Δ strains (Supplementary **Table S4**). This at first sounds conflicting with respect to the function of the deleted genes, as Sir2p is a deacetylase and Gcn5p is an acetyl transferase. However, Sir2p orthologs in human (SIRT3) and *S. enterica* (cobB) have been shown to be essential for acetyl-CoA synthetase functionality by deleting an inhibitory acetylation site (Schwer et al., 2006; Starai et al., 2002), and may thus have impact on availability of acetyl-CoA concentration and acetylation status also in *S. cerevisiae*. Working along this hypothesis, a deletion of *SIR2* would reduce available acetyl-CoA and protein acetylation level, which could help to explain the reduction in Fas2p acetylation seen in *sir2*Δ. With regard to the increased level of acetylation observed in strain *gcn5*Δ, this could possibly be explained with a reduction in histone acetylation, which could reduce expression of *ACC1*, which is dependent on histone acetylation (Galdieri et al., 2014), leading to increased acetyl-CoA and protein acetylation levels. However, as the genes investigated in the study by Kumar et al. (unpublished) are global regulators of metabolism, a consideration of FA synthesis ability of the mutants would not generate information of value for this study as it could be influenced by multiple reasons.

When discussing the potential influence of regulatory acetylations of FAS, it is worth taking the confirmed regulatory acetylation of enzymes located at the acetyl-CoA branchpoint in other species into consideration. For example, the negative regulation of Acs2 in human (Starai et al., 2002) and *Salmonella enterica* Acs (Starai et al., 2005) is a type of feed-back inhibition, as its product acetyl-CoA is involved in repressing Acs-function by donating the acetyl-groups. In line with this, it is tempting to consider that FAS acetylation could be a response of substrate-activation, which however is in disagreement with the fact that human FAS recently was found to be destabilized by acetylations, promoting ubiquitinylation and proteasomal degradation (Lin et al., 2016). It should, however, be highlighted, that human FAS is encoded as a single protein which assembles as homodimers and exhibits several important differences in structure and function compared to *Sc*FAS (Maier et al., 2008). For example, it contains a single enzymatic domain which catalyzes both acetyl and malonyl transfer and carries a thioesterase domain which releases FFAs as final products. Thus, one should be careful to draw conclusions about *Sc*FAS-regulation based on findings related to its human counterpart.

As already mentioned, acetyl-CoA concentration has been shown to be at its highest level in exponential phase when glucose is not limiting, to decrease at the diauxic shift and to remain low throughout the ethanol phase and the stationary phase (Cai et al., 2011). The gene of the proposed rate-limiting enzyme in FA biosynthesis, acetyl-CoA carboxylase (*ACC1*), is transcriptionally upregulated by increased acetyl-CoA levels via histone acetylation (Galdieri et al., 2014), and should during glucose consumption be marginally inhibited by Snf1p, which is subject to glucose repression (Kayikci and Nielsen, 2015). Based on these observations, one would assume that the phase of glucose consumption is particularly well suited for FA biosynthesis. However, yeast appears to have a greater capacity to produce FAs during growth on ethanol (Teixeira et al., 2017), which is supported by the results in this study (e.g. **Figure 3A and 3B**). A negative regulation of FAS-activity by acetylation would help to explain the lesser FFA concentrations seen in glucose phase. It should also be highlighted that 18 acetylation sites of Acc1p were detected in the acetylome data by Henriksen et al. (2012), and the potential regulatory influence of these are yet unknown. In line with this, it would be highly valuable to know if the FAS-enzyme is more highly acetylated in particular growth phases. This could possibly be addressed with the combined use of a purification step of FAS-protein from cells in different conditions and an acetyl-lysine antibody blot. Alternatively, mass-spectrometry based evaluation could be utilized.

As previously mentioned, the three K-residues selected were all evolutionary conserved among different fungal species (Jenni et al., 2007). Recently, an acetylome study was conducted in *Yarrowia lipolytica*, highlighting its FAS as being an highly acetylated enzyme (Wang et al., 2017). We performed a sequence alignment of the *S. cerevisiae* and *Y. lipolytica* FAS-sequences and found 25 out of 50 of the detected acetylated lysines in *S. cerevisiae* Fas2p to be conserved in *Y. lipolytica*, out of which four (K64 (MPT), K1092 (KS), K1446 (KS), K1821 (PPT)) also showed to be acetylated in *Y. lipolytica*. Corresponding numbers for Fas1p are 18 out of 29 acetylated lysines to be conserved, out of which four were conserved in the acetylome data set of *Y. lipolytica* (K1426 (DH), K1639 (MPT), K1753 (MPT), K2017 (MPT)). These should constitute an interesting set of acetylation sites to be investigated in potential future studies of FAS-acetylation.

## Conclusions

The results generated in this study do not allow us to determine if acetylation of K83, K173 or K1551 has a regulatory effect on Fas2p. A mutation of K173 to glutamine, suggested to be mimicking an acetylated lysine residue, slightly increased the C18/C16 ratio, but it cannot be excluded that this was due to an introduction of steric change in the catalytic chamber of FAS. It is proposed that a general strategy to evaluate impact of FAS-acetylation is adopted, as the high number of acetylation sites detected on both FAS polypeptides, Fas1p and Fas2p, makes it difficult to select the ones which have the greatest influence on activity. Alternatively, conserved acetylation sites which also have been detected in the acetylome data of other fungal species, such as *Y. lipolytica*, would have a greater likelihood of being of regulatory type.

